# Pervasive convergent evolution and extreme phenotypes define chaperone requirements of protein homeostasis

**DOI:** 10.1101/578724

**Authors:** Yasmine Draceni, Sebastian Pechmann

## Abstract

Maintaining protein homeostasis is an essential requirement for cell and organismal viability. An elaborate regulatory system within cells, the protein homeostasis network, safeguards that proteins are correctly folded and functional. At the heart of this regulatory system lies a class of specialized protein quality control enzymes called chaperones that are tasked with assisting proteins in their folding, avoiding aggregation, and degradation. Failure and decline of protein homeostasis are directly associated with conditions of aging and aging-related neurodegenerative diseases such as Alzheimer’s and Parkinson’s. However, it is not clear what tips the balance of protein homeostasis and leads to onset of aging and diseases. Here, we present a comparative genomics analysis of protein homeostasis in eukaryotes and report general principles of maintaining protein homeostasis across the eukaryotic tree of life. Expanding a previous analysis of 16 eukaryotes to 216 eukaryotic genomes, we find a strong correlation between the size of eukaryotic chaperone networks and size of the genomes that is distinct for different species kingdoms. Importantly, organisms with pronounced phenotypes clearly buck this trend. *Northobranchius furzeri*, the shortest-lived vertebrate and widely used model for fragile protein homeostasis is found to be chaperone limited. *Heterocephalus glaber* as the longest-lived rodent thus especially robust organism is characterized by above average numbers of chaperones. Our work thus indicates that the balance in protein homeostasis may be a key variable in explaining organismal robustness. Finally, our work provides an elegant example of harnessing the power of evolution and comparative genomics to address fundamental open questions in biology with direct relevance to human diseases.

## Introduction

Keeping one’s proteome properly folded and functional through varying conditions and stresses is of critical importance for cellular and organismal survival (1). Conversely, the decline of the cell’s capacity to keep its proteins in their correct shape, i.e. to maintain protein homeostasis, is a central hallmark of aging and onset of aging-associated diseases (2). Naturally, there is intense interest to better understand the principles of successful protein homeostasis as well as the origins of its failures. Cells maintain protein homeostasis through a complex regulatory network that integrates protein synthesis, folding, degradation, and trafficking pathways (3). The central players of the protein homeostasis network are a class of specialized protein quality control enzymes called chaperones (4, 5). Chaperones assist proteins in their folding, protect them from aberrant aggregation while promoting functional assembly, and, if needed, sequester and target them for degradation (4).

Eukaryotic genomes often contain more than ∼50-300 different chaperone genes that classify into distinct families based on the structure and function of the encoded proteins. As the best-characterized heat shock protein, Hsp90 type chaperones are the main stress responders that stabilize and refold partially unfolded and stress-denatured proteins (6, 7). Hsp90 chaperones interact with large and diverse sets of client proteins (6), are highly expressed, and usually encoded by only few genes (8) that are highly activated under unfavorable conditions (9). Conversely, the family of Hsp70 chaperones are key enzymes in determining the fate of newly made proteins (10): Hsp70 assist in the *de novo* folding of nascent polypeptides (11), their translocation, and disaggregation (12). Many Hsp70s are thus transcriptionally coupled to protein biosynthesis (9). Hsp40 chaperones primarily act as co-chaperones and nucleotide exchange factors to Hsp70s, thus providing specificity in guiding protein folding, assembly, disassembly, and translocation (13-15). Small heat shock proteins (sHsp or Hsp20) stabilize partially denatured proteins and prevent their aggregation (16), while enzymes from the Hsp100 chaperone family are primarily involved in disaggregation and proteolysis of proteins (17, 18). Hsp60s encode protein subunits of eukaryotic chaperonins, which are heteromeric protein complexes that provide a fully enclosed protective cavity for the folding of select proteins (17, 18). Together with their individual functional specializations, the ensemble of chaperones in the cell provides a powerful network to control the quality of proteins.

Chaperones generally bind to insoluble, sticky proteins that are at risk of aggregation (19). However, unlike in bacteria where each chaperone family is represented by only one or very few genes, chaperone genes in higher eukaryotes markedly increase in numbers. This is particularly true for chaperones of the Hsp70 and Hsp40 families that can be found with strongly increasing diversity in eukaryotes (15). The reasons why so many different types of chaperones are necessary to keep the proteome folded is far from understood. In specifically interacting with select pools of client proteins, chaperones are likely key regulatory elements of cellular networks (20). Moreover, because both biosynthesis and maintenance as well as the ATP-dependent activity of chaperones comprise substantial energetic costs, the increasing diversity in eukaryotic chaperone families has likely arisen under selection and directly reflects the requirements of managing increasingly complex proteomes (21).

To this end, the analysis of dynamically regulated proteomes in response to stress (22) as well as the comparative analysis of related organisms (23) offer the opportunity to gain fundamental insights into how the balance between the size and composition of the chaperone network as functional core of cellular protein homeostasis and the composition of the proteome is maintained (24). For instance, a general correlation between the size of the chaperone network and number of protein coding genes could be observed for 16 eukaryotic genomes (23). Malignant tumor growth in cancers is aberrantly balanced by dynamically adjusted chaperone expression profiles (25). And even viral infections impose specific requirements on the host cell chaperone network to sustain their replication (26-28).

Notably, chaperones are some of the most ancient and evolutionarily conserved components of the cell. Accordingly, evolutionary approaches offer a great opportunity to learn more about the functioning and organization of chaperone networks and cellular protein homeostasis (29-32). For instance, the evolutionary history of the Hsp90 family suggests multiple independent duplication and gene loss events (8), possibly reflecting a perpetual challenge to stay in balance with an equally evolving proteome. In turn, the expansion of proteomes (33) and encoded protein networks (34, 35) themselves is promoted by chaperones. Within proteomes, a dominant evolutionary force is given by selection to avoid protein aggregation (36). This manifests in diverse strategies to mitigate the risk of aggregation through aggregation resistant sequences (37, 38) and evolutionary design principles that protect proteins from aggregation (39, 40), strategies to temporarily sequester aggregation prone proteins (41), or increased protein quality control for aggregation prone proteins (11, 42, 43). The combinatorics of these diverse strategies hints at the complexity of understanding protein homeostasis.

Evolution offers striking examples of innovation to counter varying challenges, conditions, or constraints. In particular, the study of extreme phenotypes offers fascinating insights into the trade-offs underlying organismal fitness. At the protein level, thermophilic organisms counter the challenge of thermodynamic protein stability at elevated temperatures with reduced surface hydrophobicity to avoid aggregation yet an increase in buried sequence hydrophobicity to stabilize their protein structural core (44, 45). Similarly, specifically evolved ice-binding proteins allow species to thrive at sub-zero temperatures (46). At the cellular level, the *Dictyostelium discoideum* proteome with an extremely high number of aggregation-prone prion domains is balanced by an increase in Hsp100 disaggregase capacity (47). Recently established powerful model systems for fragile or especially robust protein homeostasis include *Northobranchius furzeri* (48, 49), the African Killifish, and *Heterocephalus glaber* (50), the naked mole rat. *N.furzeri* is the shortest-lived vertebrate. Within a life expectancy of only 4 – 6 months (48), *N.furzeri* progresses through many of the hallmarks associated with human aging including pronounced loss of protein homeostasis. The naked mole rat *H.glaber* in turn is the longest living rodent with a life expectancy of over 30 years (50). This stands in stark contrast to strongly related rodents such as *Mus musculus* that only live for around 3 years (50). In both cases the genetic origins of the comparably short and long lifespans respectively are far from understood. However, in all cases aging is characterized by the loss of protein homeostasis (51) and the accumulation of misfolded and aberrantly aggregated proteins (52), underlining the importance of better understanding how cells keep their proteomes in balance.

Here, we present a comparative genomics analysis of protein homeostasis across the eukaryotic tree of life. By significantly expanding an initial review of 16 eukaryotic genomes (23) to the quantitative analysis of 216 eukaryotes, we report general principles of keeping the balance of successful protein homeostasis. A strong correlation between the size of eukaryotic chaperone networks and size of the genomes suggests pervasive convergent evolution that follows slightly distinct trends for different kingdoms. *N. furzeri* as widely used model for fragile protein homeostasis is found to be chaperone limited, while *H.glaber* as especially robust organism is characterized by above average numbers of chaperones. Our work thus indicates that the balance in protein homeostasis may be a key variable in explaining organismal robustness.

## Results

To investigate principles of the evolution of protein homeostasis in eukaryotes, we first sought to collect a comprehensive high-quality dataset of annotated proteomes that we could analyze in the context of their phylogenetic relationships. Phylogenetic models across a highly diverse set of species are best computed based on a single marker (53), in eukaryotes usually the highly conserved 18S ribosomal RNA (rRNA) gene. Of note, the assembly and annotation state of available genomic data differs widely. Despite its central importance for phylogenetic analyses, the rRNA locus is not always annotated in early draft genomes, likely due to difficulties with mapping the many repetitive sequence elements of the intra-genic linker regions (54). Therefore, we tested different approaches to efficiently identify 18S rRNA sequences in sequenced genomes: the popular heuristic sequence similarity search algorithm BLAST (55) that is fast but not very accurate, and the hidden Markov model (HMM) based algorithm RNAmmer (56) that is very accurate but slow. Finally, to circumvent these trade-offs, we developed a pipeline based on available algorithms to readily identify 18S rRNA sequences in genomes (Fig. 1A). Specifically, we created artificial 50nt-long short reads from the annotated and validated rRNA sequences of the model organisms *C.elegans, D.melanogaster, H.sapiens, M.musculus*, and *S.cerevisiae*. that were aligned with HISTA2 (57) to the target genome to identify the candidate rRNA locus that contains the 18S rRNA sequence (*see Methods*). RNAmmer (56) was subsequently used to extract the exact 18S rRNA sequence from the candidate rRNA locus.

**Figure 1.**
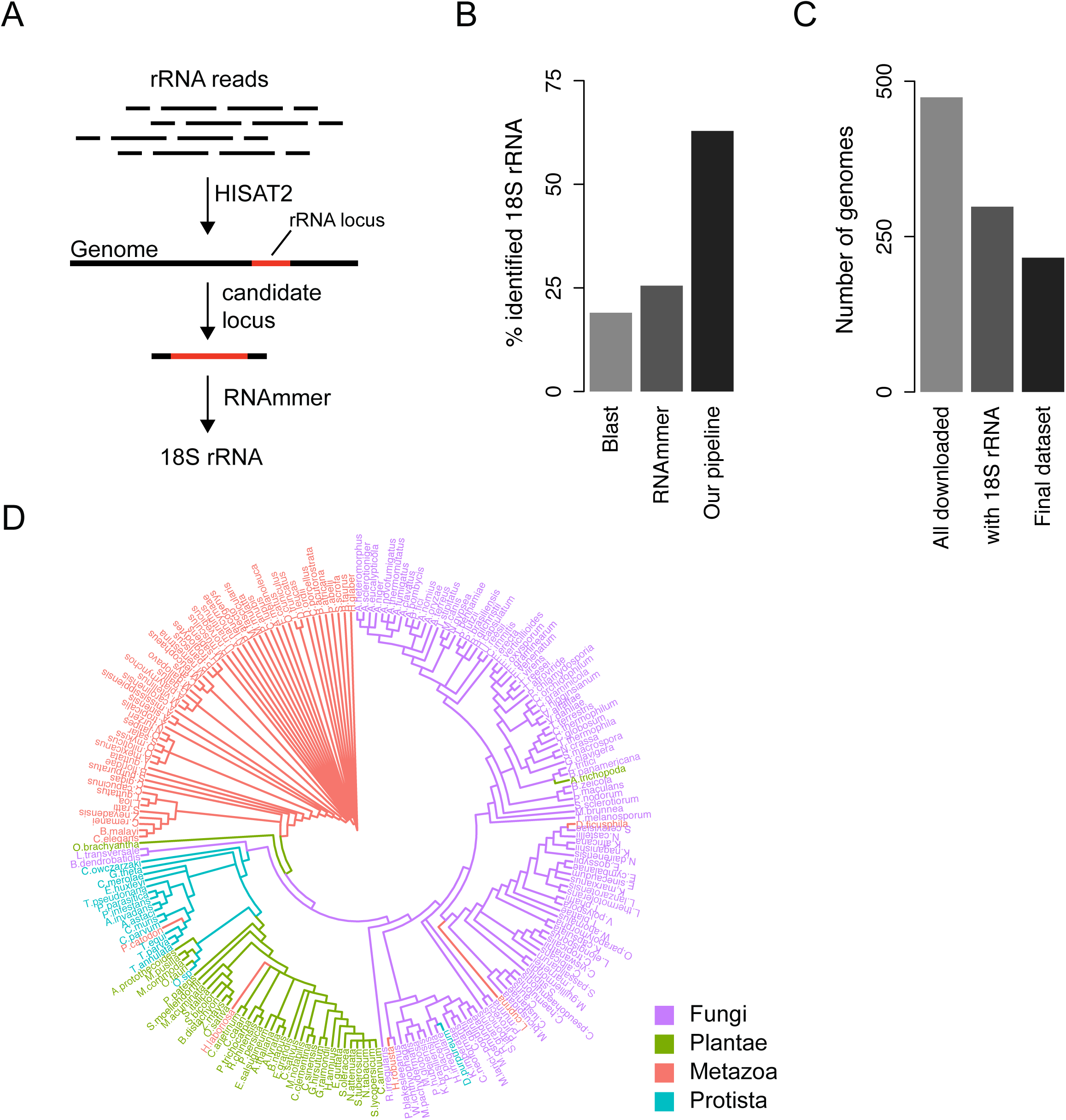
Phylogenetic classification of eukaryotes. **A** Pipeline for the efficient identification of 18S rRNA sequences in genomes based on aligning short rRNA reads with HISTA2 and RNAmmer. Mapping of synthetic rRNA reads is used to find the best candidate locus, followed by exact 18S rRNA identification with RNAmmer. **B** Benchmarking the identification of rRNA. The developed pipeline improves the efficient identification of 18S rRNA sequences in genomes (Table S1). **C** Data quality control. All genomes with genome assemblies in the RefSeq database and corresponding validated proteomes in the Uniprot database were downloaded. Not all genome assemblies include the 18S rRNA locus, and few additional organisms were excluded from further analyses, either because their 18S rRNA sequence contained ‘N’ characters, or because the 18S sequences introduced above average numbers of gaps into the multiple sequence alignment (*see Methods*, Table S2). **D** Eukaryotic tree of life coloured by species kingdoms. Strong clustering of species within their kingdoms supports a good phylogenetic model.

We were able to identify 18S rRNA sequences by BLAST in only 19%, and by RNAmmer in only 25% of the eukaryotic genomes from the RefSeq databse (Fig. 1B; Table S1). Moreover, BLAST yielded many false positive hits that upon closer inspection did not align well with validate 18S rRNA sequences. We concluded that BLAST was not an appropriate tool to discover rRNA sequences. The software RNAmmer is highly accurate in the identification of rRNA but too slow to efficiently scan full genomes. Therefore, we could only obtain 18S rRNA with RNAmmer from genomes where the rRNA locus had already been annotated. With our strategy to use the short-read aligner HISAT2 and synthetic rRNA reads from a mix of source organisms to map the best candidate rRNA locus followed by the accurate identification of the 18S rRNA with RNAmmer we could expand our dataset to 62% of the initially downloaded genomes (Fig. 1B; Table S1). The missing rRNA sequences in the remaining genomes likely suffer from challenge to resolve the repetitive sequences in the rRNA cassette in early genome assembly states. Our results suggest that the combination short-read alignment and RNAmmer can be a very efficient manner to identify rRNA sequences in genomes. Additional conservative data quality control (*see Methods*; Fig. 1C; Fig. S1A, B) resulted in a set of 216 eukaryotes spanning Fungi, Protista, Metazoa, and Plantae, i.e. all four eukaryotic kingdoms of life, for which we computed a phylogenetic tree (Fig. 1D). The obtained phylogenetic model was supported by a good *maxdiff* = 0.27 between three independent trees and found in very good agreement with taxonomy annotations (Fig. 1D). Moreover, the consistent clustering of species within their kingdoms quantified to clear differences in shorter pairwise evolutionary distances between species within their cluster compared to longer distances between species from differetn kingdoms (Fig. S1C). Protists were found the least separated from the other kingdoms, which may also be the result of low representation in our dataset. In contrast, both animals and plants partitioned into well-defined and distinct clusters (Fig. 1D).

We next wanted to better understand principles of the evolution of chaperone networks as core components of the protein homeostasis network across this eukaryotic phylogeny. All occurrences of nuclear genome encoded chaperones of the Hsp20, Hsp40, Hsp60, Hsp70, Hsp90, and Hsp100 families were identified (*see Methods*) and mapped onto the phylogenetic tree (Fig. 2). Strikingly, patterns of the counts of chaperone genes directly correlate with similarities and divisions in the phylogeny. Increasing organism complexity is accompanied by increasing diversity in all chaperone families (Fig. 2). Especially visible is the clear transition from animals to the polyploid plant genomes (Fig. 2). The shift to larger chaperone networks for more complex genomes is noticeable for all chaperone families, but especially standing out are increased counts for Hsp20, Hsp40, and Hsp70 type chaperones (Fig. S2). Within the different kingdoms, plants show by far the largest variance in the composition of their chaperone networks (Fig. 2, S2). While the composition of eukaryotic chaperone networks appears in general to be strongly correlated, individual species are clearly standing out by above or below average numbers in individual chaperone families (Fig. 2). Taken together, our analysis of the chaperone networks across eukaryotes suggests fundamentally conserved similarities with pronounced exceptions.

**Figure 2.**
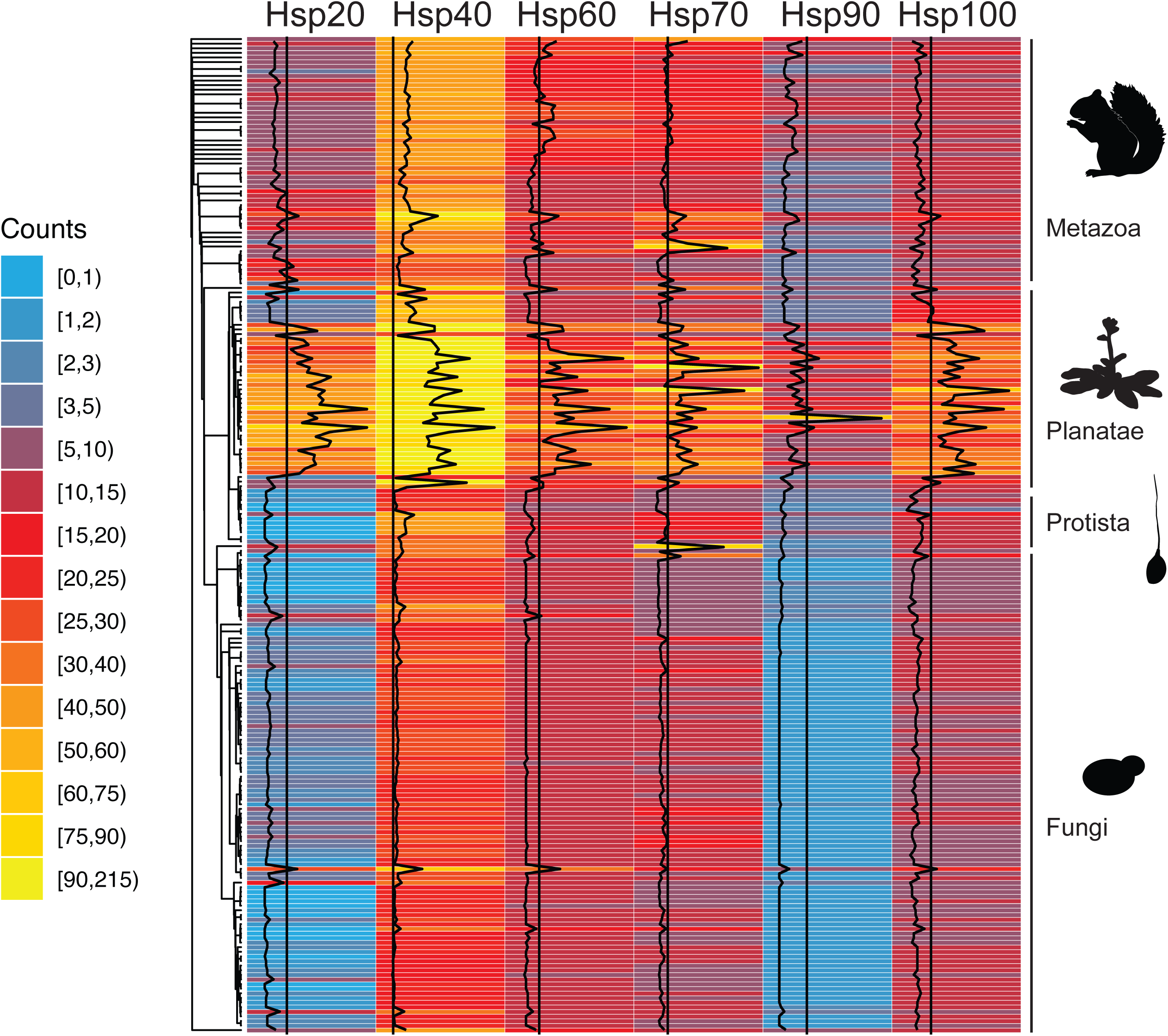
Evolution of chaperone networks. The counts of nuclear-encoded Hsp20, Hsp40, Hsp60, Hsp70, Hsp90, and Hsp100 chaperone genes from 216 eukaryotic genomes are visualized as heatmap. Black lines indicate the absolute numbers of chaperone genes relative to their median across all species for each chaperone family.

We next systematically evaluated these observed (dis)similarities in the composition of eukaryotic chaperone networks as function of their evolutionary distance. Pairwise chaperone network similarity was quantified by the correlation coefficient between the chaperone counts in the six protein families, and compared to the evolutionary distances obtained from the branch lengths of the phylogenetic tree. To highlight organisms that differ the most from the rest, we computed for each species the average of its pairwise correlation coefficients to the chaperone profiles of all others species, and similarly the average of the pairwise evolutionary distances to all other species (*see Methods*). Focusing on the most populated clusters, i.e. the fungi (Fungi), animals (Metazoa), and plants (Plantae), we observed a general strong correlation of the composition of the chaperone networks independent of average evolutionary distance (Fig. 3A). In the Fungi and Plantae groups almost all species had average correlation coefficients above 0.9, thus suggesting that the chaperone networks in eukaryotes are generally composed of similar relative numbers of Hsp20, Hsp40, Hsp60, Hsp70, Hsp90, and Hsp100 type chaperones. Because these different chaperones perform overlapping yet specialized functions it is not surprising that all of them are proportionally present. Moreover, the observation that the generally very high similarity in chaperone profiles across eukaryotes is found to be independent of evolutionary distance (Fig. 3A) indicates a fundamental selective pressure on proteome composition, likely through convergent evolution.

**Figure 3.**
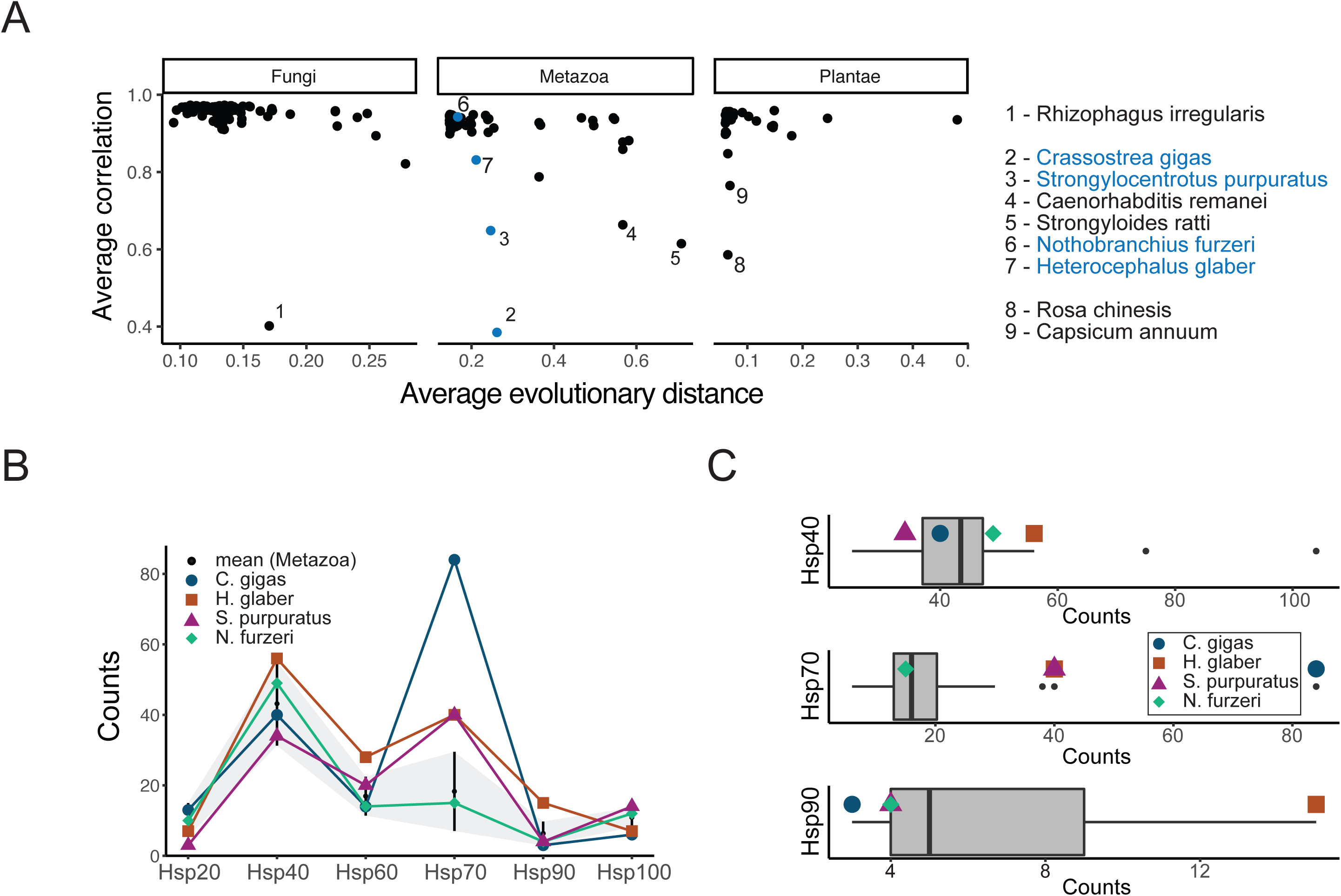
Conservation and diversity of in the composition of chaperone networks. **A** Similarity of the composition of chaperone networks within phylogenetic clusters as function of evolutionary distance. Average evolutionary distance denotes the average of pairwise evolutionary distances computed for the branch lengths. Average correlation coefficients represent the average of pairwise correlation coefficients between the composition of chaperone networks of a query species to the other species. **B** Composition of the chaperone networks of four exemplary and noteworthy species that are characterized by known extreme phenotypes, namely the shortest-lived vertebrate *N.furzeri*, the longest lived rodent *H.glaber*, the pacific oyster *C.gigas*, and the sea urchin *S.purpuratus.* As reference are shown the distributions of the chaperone counts in all Metazoa (black dots) +/- standard deviation (grey area). **C** Distribution of the number of chaperones of different families across the animal kingdom. The four species are highlighted in their extreme positions relative to all other animals analyzed.

The fungus with the lowest average correlation coefficient was *Rizophagus irregularis*, a symbiotic fungus used in soil agriculture that belongs to a different taxonomy division than the rest of the fungi in our data. The plants with the lowest average correlation coefficient in our dataset were *Rosa chinesis* and *Capsicum annuum*, two highly domesticated plants (Fig. 3A). In contrast, the cluster of animal species in our dataset exhibited clearly more diversity. Specifically, the Pacific oyster *Crassostrea gigas* was found to have the most dissimilar chaperone profile with an average correlation of only 0.38, followed by the sea urchin *Strongylocentrotus purpuratus* (average correlation = 0.65) despite short average evolutionary distances. We decided to analyse more in-depth the chaperone networks of these two dissimilar animals found by this analysis, *C.gigas* and *S.purpuratus*, together with two organisms of known phenotype of interest, *N.furzeri* and *H.glaber* (Fig. 3B, C). Upon closer inspection it became apparent that *C.gigas* had an extraordinary high count of Hsp70 chaperones at over 80 (58) while the average across the studied animal genomes was only around 15 (Fig. 3B,C). However, *C.gigas* only has 6 Hsp100 (Fig. S3). The sea urchin *S.purpuratus* was equally characterized by a high number of over 40 Hsp70 chaperones (Fig. 3B,C) but also one of the highest observed count of Hsp100 (Fig. S3). Strikingly, the naked mole rat *H.glaber* displayed one of the highest counts of Hsp40 chaperones and ranked amongst the highest counts of both Hsp70 and Hsp90 chaperones of all animals studied (Fig. 3B,bC). In contrast, the African killifish *N.furzeri* stood out for one of the lowest observed counts of Hsp90 chaperones among the group of animals (Fig. 3B, C). Taken together, all examples that represent extreme phenotypes were indeed also characterized by extreme counts of specific chaperone types as compared within the group of animal genomes analyzed.

Because chaperones preferentially bind to aggregation prone proteins, we next quantified the aggregation propensity of the proteomes across the phylogeny (Fig. 4A). The distributions of per-protein predicted aggregation propensity were represented by the characteristic median and lower and upper percentiles and clearly correlated well for many closely related species (Fig. 4A). On average, fungi and animals showed a higher aggregation scores than plants (Fig. 4B). The combination of both higher chaperone counts and lower aggregation scores in plants hints at the complexities of their life styles. Remarkably, the median aggregation score of the proteomes of *C.gigas, S.purpuratus, H.glaber*, and *N.fuzeri* fell right on the center of the corresponding distribution of all animals studied, or even placed on the lower end of the 25% percentile (Fig. 4C). This suggests that these organisms do not possess proteomes with elevated aggregation scores.

**Figure 4.**
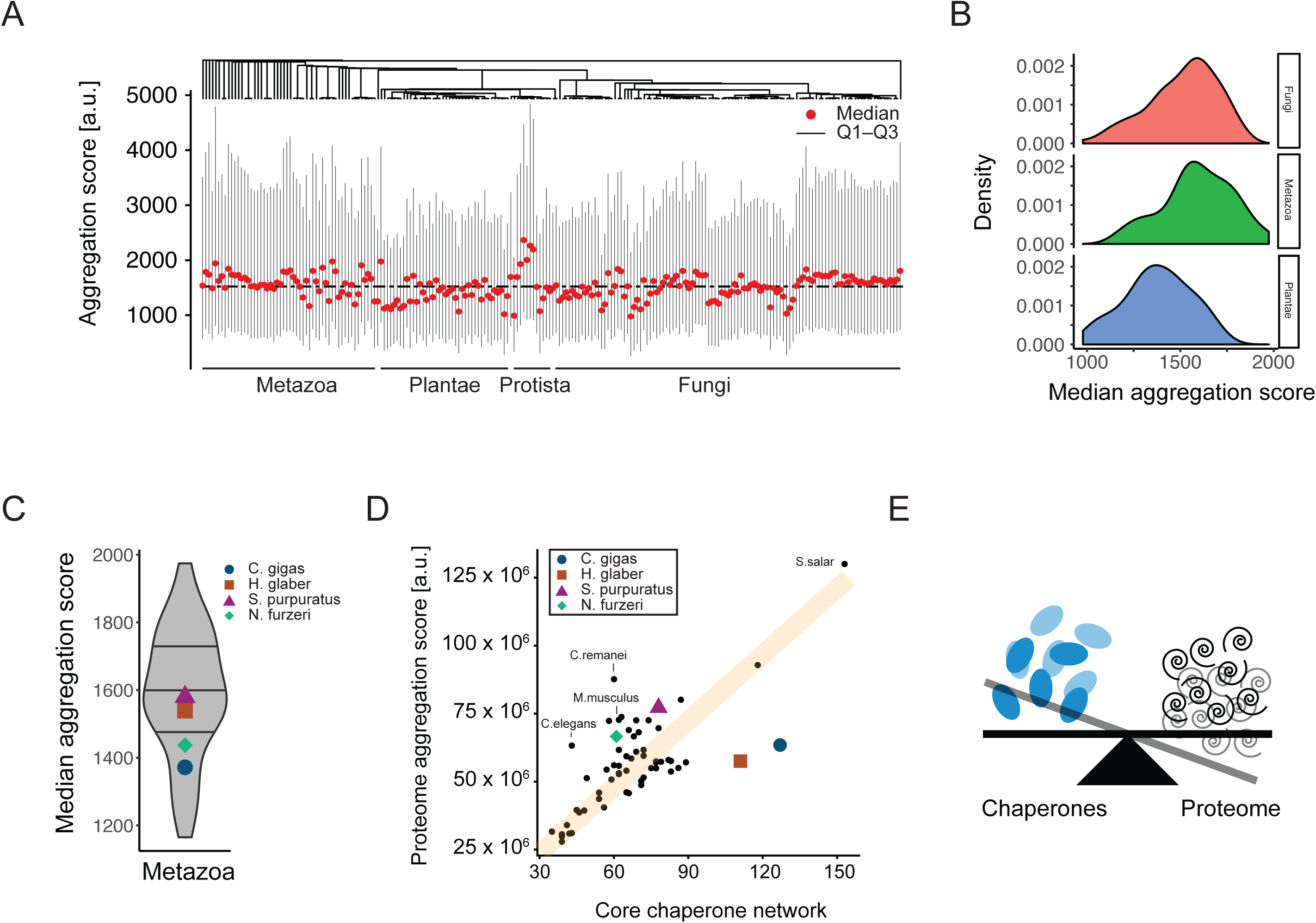
Keeping the proteome in balance. **A** Distribution of predicted protein aggregation propensities in proteomes across the phylogeny. Per-protein predicted aggregation scores are represented by median and 25% (Q1) and 75% (Q3) percentiles for each species. **B** Distributions of the representative proteome median aggregation scores for the four clusters of species kingdoms on the phylogenetic tree, namely Fungi, Protista, Metazoa, and Plantae. **C** Distribution of median aggregation scores across the animal kingdom. The median aggregation scores of *C.gigas, S. purpuratus, N.furzeri* and *H.glaber* are highlighted. **D** Relationship between the size of chaperone networks and proteome aggregation propensities. The protome aggregation score was computed as the sum of the per-protein predicted aggregation scores, thus reflecting the size of a proteome weighted by the predicted propensity of its proteins to aggregate. The trend line (orange line) is indicative and not a linear fit to the data. **E** Chaperone networks and proteome co-evolve to maintain balance in protein homeostasis while extreme phenotypes correlate with extreme chaperone counts.

Biological organisms are most beautifully complex systems, and complex phenotypes can rarely be explained by individual observables. To understand how species may balance their proteome with costly chaperone capacity, we next compared the size of the chaperone networks in individual species with the overall aggregation score of their proteomes (Fig. 4D). Specifically, we first considered Hsp40, Hsp70, and Hsp90 as the core chaperone network as these are the main chaperones for initial folding, stress response, and disaggregation. A proteome aggregation score was computed as the sum of the individual predicted protein aggregation propensities, thus reflecting the size of the proteome weighted by its propensity to aggregate. Remarkably, we observed a general and very strong linear correlation between the size of the core chaperone network and the proteome aggregation score (Fig. 4D) that is much more pronounced that what had previously been reported for a subset of 16 eukaryotes (23). Several organisms clearly diverted from this trend including the ones previously identified. The naked mole rat *H.glaber* was found far to the right of this general trend, suggesting an excess of chaperones compared to the proteome aggregation propensity and relative to the other animals (Fig. 4D). This observation matches very well the remarkable robustness and longevity of *H.glaber.* The Pacific oyster *C.gigas* could also be found far to the right of the general trend (Fig. 4D). In contrast, the African killifish *N.fuzeri* placed on the left side of the general correlation (Fig. 4D), reflective of its known fragile protein homeostasis. Similarly, *Mus musculus* as one of the shortest-lived mammals equally placed clearly to the left side of this correlation, as did the nematodes *C.elegans* and *C.remanei* and (Fig. 4D). The sea urchin *S.purpuratus* however fell onto the general trend, suggesting that its expanded chaperone network merely compensates a more aggregation prone proteome and other putative requirements (Fig. 4D). Importantly, the same observations could be made when considering all chaperone families (Fig. S4A). Moreover, the analysis of our complete dataset across four kingdoms revealed that in individual species kingdoms follow slightly different trends with a decreasing slope from fungi and protists to animals and finally plants (Fig. S4B). Taken together, our results reveal fundamental principles of keeping protein homeostasis in balance that explain known cases of particularly weak or robust protein homeostasis.

Next to these global trends in the balance between chaperones and proteome, our observations match many fascinating facets of species-specific constraints and adaptations. The sea urchin *S.purpuratus* is an important model organism for developmental biology and characterized by a remarkable lifespan of over a century as well as high fecundity (59, 60). With a high degree of polymorphisms in its population (60) and strong defense systems (61), *S.purpuratus* contained one of the largest counts of Hsp100 chaperones of all animals analyzed. The proteolytic activity of Hsp100s may assist the complex and sophisticated innate immune system (62) in defending against a plethora of stressors, as well as the biosynthesis and assembly of its pronounced spikes. Another maritime organisms, the Pacific oyster *C.gigas* has evolved unusually high numbers of chaperones to adapt to harsh living conditions and the specific biosynthesis requirements of making the hard oyster shells (58). Also living in the highly stressful intertidal zone, the genome is characterized by a remarkable expansion of Hsp70s (58). Moreover, the oyster shell is formed by a diverse array of matrix proteins through complex assembly and modification processes (58). Hsp70 chaperones frequently function as assembly factors. Thus, next to an extraordinary capability to adapt to stress conditions (63, 64), the large number of Hsp70s may play a crucial role in shell formation. With respect to longevity, the most striking observation from our analyses is the opposing numbers of Hsp90 proteins in *N.furzeri* and *H.glaber*. The short-lived *N.furzeri* has the lowest and the long-lived *H.glaber* the highest number of Hsp90 genes in the group of Animalia. While both proteomes are likely well balanced under normal conditions, the ability to adapt to stress conditions is critical for long-term survival. While *N.furzeri* has strong capacity in the Hsp20 and Hsp40 systems, Hsp70 is equally only present in relatively few copies. The naked mole rate *H.glaber* is often compared to *M.musculus* for their stark contrast in longevity (65) and here equally stands out the much higher numbers of Hsp70 and Hsp90 genes in the longer-lived *H.glaber* (Fig. 4D). Another short-lived model organisms for aging research, the nematode *C.elegans*, is equally found to be chaperone limited (Fig. 4D). The individual living conditions and phenotypes of different organisms are as complex as the protein homeostasis network, that will ultimately have to be understood at a much more detailed level. However, our comparative genomics analyses reveal fundamental trends of how organisms keep their proteome in balance, and how diversion from this trend correlates with known extreme phenotypes.

## Discussion

We have performed a comparative genomics analysis of protein homeostasis in 216 eukaryotes. Our results suggest strong convergent evolution to maintain the overall composition of eukaryotic chaperone networks across the eukaryotic kingdoms of life. Strikingly, those organisms that divert from the general chaperone profile are often accompanied by specific phenotypic constraints. Some of the shortest-lived species have the lowest numbers of Hsp90 genes, the main stress response chaperone, while especially robust organisms are characterized by high Hsp90 chaperone counts. Our finding that the composition of chaperone networks with proportional numbers of Hsp20, Hsp40, Hsp60, Hsp70, Hsp90, and Hsp100 is generally very stable and independent of evolutionary distance underlines that this evolutionarily ancient machinery strongly co-evolved with the corresponding proteomes. In turn, the finding that the cases with clearly different chaperone networks respond to special circumstances offers a fascinating window into the comparative genomics of protein homeostasis.

A general short-coming of our analysis lies in the inference from the number of nuclear encoded chaperone genes to their protein folding capacity in the cell, which equally strongly depends on their expression profiles. Nonetheless, from an engineering and systems perspective there are fundamental differences between having few loci that are highly expressed and many gene copies, even if both result in similar protein levels. As organisms age and accumulate mutations, the biosynthesis of chaperones themselves may be negatively affected. By distributing the genetic information across several loci, the system becomes more redundant thus robust (66). Equally importantly, diversity in chaperones likely yields a gain in specificity and fidelity in regulation, thus affording more control over adaptive responses to stress. Our results and the results by others suggest that this is a fundamental constraint on managing more complex proteomes of higher eukaryotes, but likely similarly affects organismal fitness and longevity.

As much as our work highlights the power of comparative genomics to shed light onto fundamental open questions in biology, as much does our work reveal current challenges and opportunities for further improvement. Large-scale comparative genomics analyses are strongly dependent on the quality and progress of genome assemblies and annotations, on their own formidable challenges. While final numbers of genes including chaperone encoding genes likely continue to change with progressing assembly and annotation, our numbers are in overall very good agreement with previously published analyses of chaperone systems.

Finally, protein homeostasis is achieved through a truly complex regulatory network and protein folding under chaperone assistance only comprises one central aspect of it (67). Many parts of this regulatory system are tightly controlled, and even RNA molecules are chaperoned (68). An alternative means to remove misfolded and aggregated proteins is provided by powerful degradation systems whose malfunctioning is equally implicated in organismal aging and disease (69). The continued explosion of sequenced genomes without doubt will enable to significantly expand evolutionary and comparative genomics efforts to learn about protein homeostasis.

## Methods

### Code availability

Computer code to reproduce all results and analyses is available at github.com/pechmannlab/chapevo

### Data sources

*W*e downloaded all eukaryotic genomes from the RefSeq database (70) that also had corresponding validated proteomes available in the Uniprot database (71). This yielded a set of 472 eukaryotic genomes and corresponding proteomes. From RefSeq, both annotated RNA sequences and whole genome FASTA files were downloaded. Draft genome assemblies of *Heterocephalus glaber* (50) (“naked mole rat”) and *Nothobranchius furzeri* (48, 49) (“African killifish”) were downloaded individually. The corresponding protein sequences for each organism were retrieved from the Uniprot database using the provided FASTA files without isoforms, and in the case of *N.furzeri* from the species-specific database (http://nfingb.leibniz-fli.de/) and filtered for one isoform per gene based on gene IDs.

### Identification of chaperones

We used the expert-curated profile hidden Markov models (HMMs) from the Heat Shock Protein Information Resource database (72) and the *hmmsearch* module of the Hmmer (73) software to identify Hsp20, Hsp40, Hsp60, Hsp70, Hsp90 and Hsp100 chaperones in sequenced and annotated proteomes. All hits of *hmmsearch* with e-values below 10^-5^ were considered.

### Identification of 18S rRNA

Three approaches were tested to efficiently find the 18S rRNA gene in genomes. First, annotated 18S rRNA sequences from the well-characterized model organisms *H. sapiens, A.thaliana, D.melanogaster* and *R.norvegicus* were searched with BLAST (55) against the target genome. Herein, best hits with *blastn* and e-values < 10^-6^ as widely accepted threshold for homology searches (74) were considered as putative rRNA loci. BLAST is, in generally, insufficient to accurately detect 18S rRNA genes as we obtained many false positive hits. Next, the software RNAmmer (56) was used to identify 18S rRNA sequences from genomes. RNAmmer is a highly accurate HMM-based software but too slow to search whole genomes. Therefore, 18S rRNA sequences could only be identified from genomes with already annotated rRNA loci. Last, we converted the validated 18S rRNA sequences from *C.elegans, D.melanogaster, H.sapiens, M.musculus*, and *S.cereviceae* with a sliding window of length 50 into artificial short RNA reads. The choice of source organisms was arbitrary but found to yield robust results. For each target genome an alignment index was built and the short-read aligner HISAT2 (57) was used to align the mix of rRNA reads form the different species to the full target genome sequence. From the resulting SAM file, the chromosome and locus with the highest coverage was identified, and the start and end positions both extended as candidate rRNA locus. We obtained robust results for extending the candidate locus by 100, 500, or 1500nt. RNAmmer was next used to efficiently identify the coordinates and sequence of the 18S rRNA within this candidate locus.

### Phylogeny of eukaryotes

We removed 28 18S rRNA sequences that contain “N” characters as they impede accurate evolutionary inference. To construct a high-quality sequence alignment, the structure-based SSU-align (75) program was used as alignment tool. Next, a phylogenetic tree was constructed with PhyloBayes (76, 77) using the CAT/GTR model and three independent Markov Chain Monte Carlo (MCMC) chains of length 10,000. The first 1000 trees were cast-off as burn-in. Our initial tree showed a largest discrepancy observed across all bipartitions of *maxdiff* = 0.534, thus suggesting under-sampling. Specifically, upon manual inspection of the 18S rRNA alignment it was clear that only few sequences caused the multiple-sequence alignment to expand by an above average number of gaps. To identify sequences that caused the largest number of gap openings, we computed pairwise sequence alignments of all sequences in our 18S rRNA dataset with SSU-align and counted the number of gaps. We considered the ‘% gaps’ as the number of gaps relative to alignment length in pairwise alignments. The ‘average % gaps’ thus represents the average percentage of gaps a sequence introduces to all other sequences in pairwise alignments. In trying to obtain the best tree, we sought to minimize the *maxdiff* while keeping as many sequences as possible. Thus, phylogenetic trees were computed upon removal of sequences exceeding six different thresholds for the ‘average % gaps’: 15%, 20%, 25%, 30%, 50%, 100%. The threshold of 15% was found to yield the best results, and sequences with a gap score of > 15% were discarded (Table S2). This yielded a dataset of 216 eukaryotic genomes. The largest discrepancy between three independent chains reduced to *maxdiff* = 0.27, thus indicating a good phylogenetic model (76).

### Taxonomy classification

Species taxonomy information was retrieved from the NCBI taxonomy database (https://www.ncbi.nlm.nih.gov/taxonomy) and assigned with the R package myTAI (77). We assigned species with missing kingdom annotations to the cluster of “Protists”.

### Aggregation scores

The propensity of proteins to aggregate was predicted from the protein amino acid sequences with the Tango (79) software. Tango predicts per-residue aggregation scores that can be summed up to per-protein scores. We considered the distribution of all predicted per-protein aggregation scores as represented through the median and characteristic 25% (Q1) and 75% (Q3) percentiles. Finally, *proteome aggregation scores* were computed as the sum of the individual protein aggregation scores, thus reflecting the size of the proteome weighted by its aggregation propensity.

### Data analysis

All data analyses were performed with custom computer code written in Python and R (github.com/pechmannlab/chapevo). Figures were prepared in R including use of the packages *ggplot2* (80), *ggtree* (81, 82), and *cowplot* (https://github.com/wilkelab/cowplot).

## Acknowledgements

We are grateful to Emmanuel Noutahi and Simon Laurin-Lemay for assistance with the PhyloBayes calculations, and the members of the Pechmann Lab for helpful discussions. This work was funded through financial support from an NSERC Discovery Grant and the Université de Montréal. Computations were in part performed on supercomputers managed by Calcul Québec and Compute Canada that are funded by the Canada Foundation for Innovation (CFI), Ministère de l’Économie, des Sciences et de l’Innovation du Québec (MESI) and le Fonds de recherche du Québec – Nature et technologies (FRQ-NT). SP holds the Canada Research Chair in Computational Systems Biology.

## Author contributions

YD developed, implemented, and carried out all computational analyses and prepared the initial manuscript and figures. SP devised and supervised the project and assisted in the preparation of the final manuscript and figures.

