## Supplement for "Pervasive convergent evolution and extreme phenotypes define chaperone requirements of protein homeostasis"

Data and code availability: github.com/pechmannlab/chapevo

This Supplement contains:

Figure S1

Figure S2

Figure S3

Figure S4

Table S1

Table S2

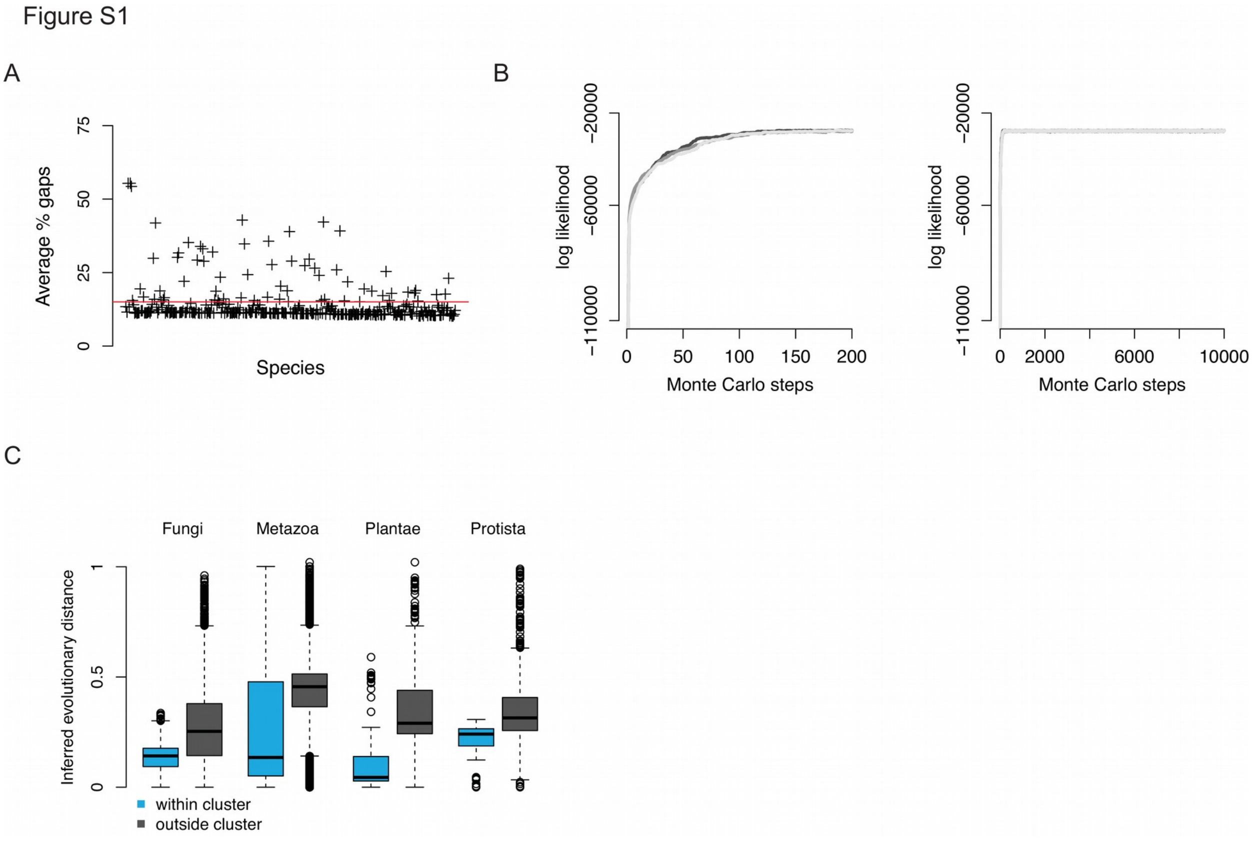


**Figure S1**. Reconstruction and validation of a phylogenetic tree of eukaryotes. **A** Identification of 18S rRNA sequences that introduce an above average number of gaps into the multiple-sequence alignment of eukaryotic 18S rRNA. For each species, the 'average % gaps' score denotes the average number of gaps relative to alignment length in pairwise sequence alignments of a query 18S sequence to the 18S sequences of all other species. The final phylogenetic tree was computed for a dataset with sequences that introduced on average at most 15% gaps (red line indicates this threshold). **B** Convergence of phylogenetic inference. Markov Chain Monte Carlo (MCMC) sampling converges quickly and reproducibly for three independent chains (left: zoom-in on first 200 steps; right: full traces) **C** Comparison of the distributions of pairwise evolutionary distances inferred from the branch lengths of the phylogenetic tree within and outside of clusters as defined by taxonomy annotations. Evolutionary distances are computed by adding up individual branch lengths with the built-in function of the ETE3 toolkit. The observed throughout shorter evolutionary distances between species from the same kingdom as compared to species outside the kingdom underline the quality of the phylogenetic model.

### Supplementary Figure 2

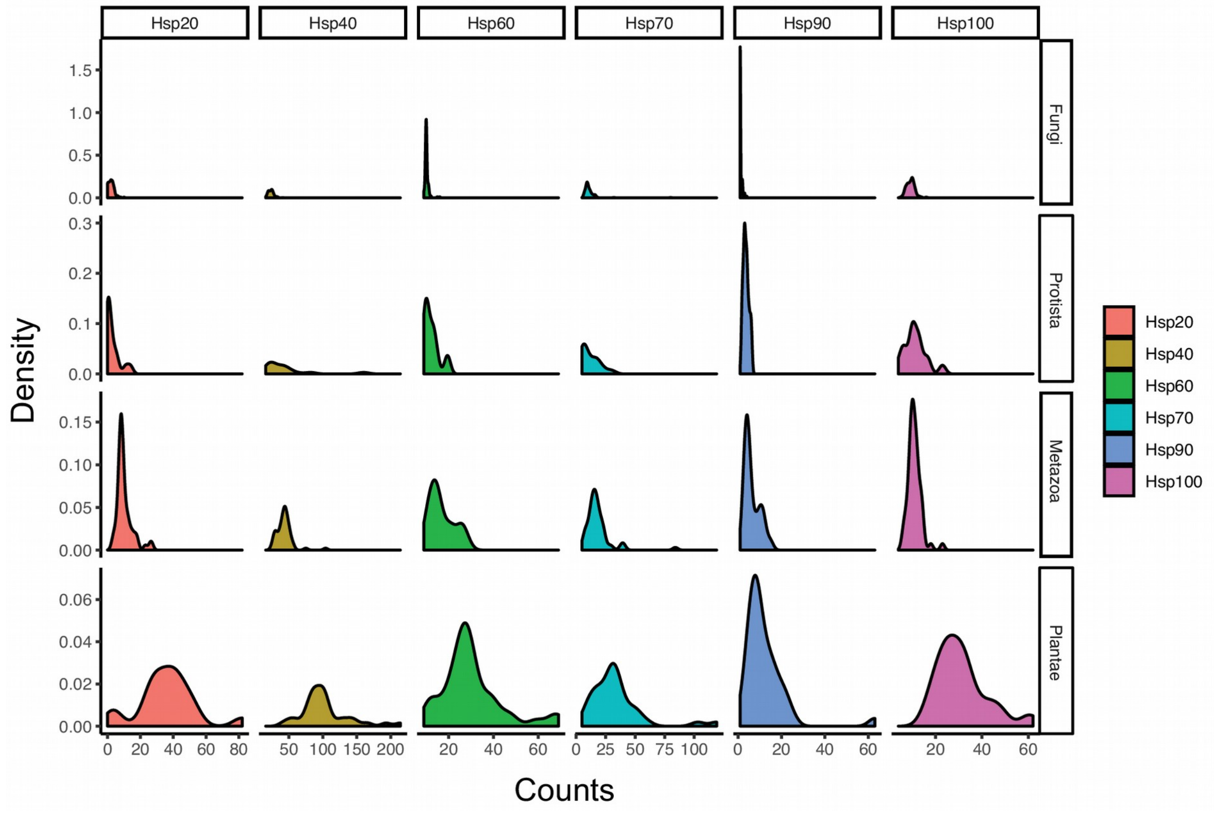


**Figure S2**. Diversity in chaperone families across the eukaryotic tree of life. Distributions of the numbers of chaperone genes in the chaperone families Hsp20, Hsp40, Hsp60, Hsp70, Hsp90, and Hsp100 are shown for the four kingdom of Fungi, Protista, Metazoa, and Plantae. Especially Hsp40 and Hsp70 type chaperones strongly increase in numbers in higher eukaryotes.

Figure S3

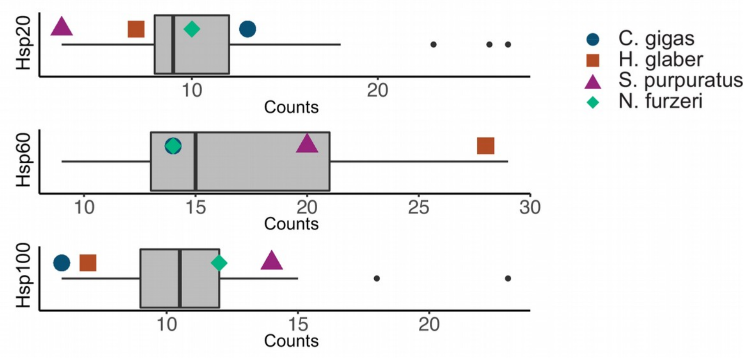


**Figure S3**. Distribution of the number of Hsp20, Hsp60, and Hsp100 chaperone genes in animal genomes. The determined counts of *C.gigas*, *H.glaber*, *S.purpuratus*, and *N.furzeri* are highlighted and often occupy extreme positions in the distributions.

Figure S4

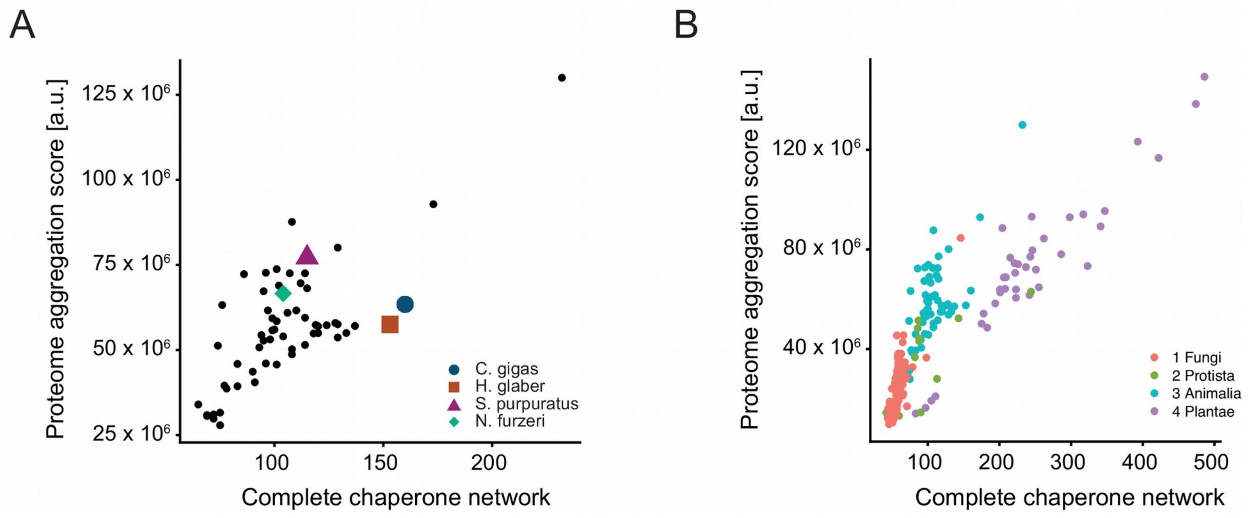


**Figure S4**. Keeping the proteome in balance. **A** Relationship between the size of the complete chaperone network and proteome aggregation propensity for animal genomes. **B** Relationship between the size of the complete chaperone network and proteome aggregation propensity for all studied genomes. The different kingdoms follow distinct trends.

| 18S rRNA | BLAST | RNAmmer | Our pipeline |
|---|---|---|---|
| **Count** | **88/472** | **119/472** | **296/472** |
| **Percentage** | **18.6** | **25.1** | **62.4** |

**Table S1**. Identification of 18S rRNA sequences in genomes. Shown are the counts of confidently obtained 18S rRNA sequences compared to the number of genomes analyzed by BLAST, RNAmmer, and our new pipeline that combines NGS rRNA read mapping with RNAmmer analysis. The two additional genomes of *N.furzeri* and *H.glaber* were analyzed separately.

| Average % gap threshold | Number of sequences | Max diff |
|---|---|---|
| **15 %** | 216 | 0.268 |
| **20 %** | 247 | 0.540 |
| **25 %** | 253 | 0.382 |
| **30 %** | 262 | 0.855 |
| **50 %** | 269 | 0.440 |
| **100 %** | 270 | 0.534 |

**Table S2**. Benchmarking phylogenetic inference with PhyloBayes. The *maxdiff* difference between independent MCMC chains is shown for datasets that were filtered at 'average % gaps' thresholds of 15%, 20%, 25%, 30%, 50%, and 100%. . A *maxdiff* < 0.3 is usually considered sufficient for a good model fit, whereas higher values suggest undersampling. The 18S sequences of 28 genomes contained "N" characters and were thus removed from our dataset.
